# Glucocerebrosidase reduces the spread of protein aggregation in a *Drosophila melanogaster* model of neurodegeneration by regulating proteins trafficked by extracellular vesicles

**DOI:** 10.1101/2020.05.15.097766

**Authors:** Kathryn A. Jewett, Ruth E. Thomas, Chi Q. Phan, Gillian Milstein, Selina Yu, Leo J. Pallanck, Marie Y. Davis

## Abstract

Abnormal protein aggregation within neurons is a key pathologic feature of Parkinson’s disease (PD). The spread of protein aggregates in the brain is associated with clinical disease progression, but how this occurs remains unclear. Mutations in the gene *glucosidase, beta acid 1* (*GBA*), which encodes the lysosomal enzyme glucocerebrosidase (GCase), are the most penetrant common genetic risk factor for PD and dementia with Lewy bodies, and also associate with faster disease progression. To explore the mechanism by which mutations in *GBA* influence pathogenesis of these diseases, we previously created a *Drosophila* model of *GBA* deficiency (*Gba1b*) that manifests neurodegeneration, motor and cognitive deficits, and accelerated protein aggregation. Proteomic analysis of *Gba1b* mutants revealed dysregulation of proteins involved in extracellular vesicle (EV) biology, and we found altered protein composition of EVs from *Gba1b* mutants. To further investigate this novel mechanism, we hypothesized that *GBA* may influence the spread of pathogenic protein aggregates throughout the brain via EVs. We found that protein aggregation is reduced cell-autonomously and non-cell-autonomously by expressing wildtype GCase in specific tissues. In particular, accumulation of insoluble ubiquitinated proteins and Ref(2)P in the brains of *Gba1b* flies are reduced by ectopic expression of GCase in muscle tissue. Neuronal expression of GCase also cell-autonomously rescued protein aggregation in brain as well as non-cell-autonomously rescued protein aggregation in muscle. Muscle-specific *GBA* expression rescued the elevated levels of EV-intrinsic proteins and Ref(2)P found in EVs from *Gba1b* flies. Genetically perturbing EV biogenesis in specific tissues in the absence of GCase revealed differential cell-autonomous effects on protein aggregation but could not replicate the non-cell-autonomous rescue observed with tissue-specific *GBA* expression. Additionally, we identified ectopically expressed GCase within isolated EVs. Together, our findings suggest that GCase deficiency mediates accelerated spread of protein aggregates between cells and tissues via dysregulated EVs, and EV-mediated trafficking of GCase may partially account for the reduction in aggregate spread.

**Author’s Summary:** Parkinson’s disease (PD) is a common neurodegenerative disease characterized by abnormal clumps of proteins (aggregates) within the brain and other tissues which can lead to cellular dysfunction and death. Mutations in the gene *GBA*, which encodes glucocerebrosidase (GCase), are the strongest genetic risk factor for PD, and are associated with faster disease progression. GCase-deficient mutant flies display features suggestive of PD including increased protein aggregation in brain and muscle. We found that restoring GCase protein in the muscle of mutant flies reduced protein aggregation in muscle and the brain, suggesting a mechanism involving interaction between tissues. Previous work indicated that *GBA* influences extracellular vesicles (EVs) – small membrane-bound structures released by cells to communicate and/or transport cargo from cell to cell. Here, we found increased aggregated proteins within EVs of mutant flies, which was reduced by restoring GCase in muscle. In addition, we found GCase within the EVs, possibly explaining how GCase in one tissue such as muscle could reduce protein aggregation in a distant tissue like the brain. Our findings suggest that GCase influences proteins within EVs, affecting the spread of protein aggregation. This may be important to understanding PD progression and could uncover new targets to slow neurodegeneration.

## Introduction

Parkinson’s disease (PD) is the most common neurodegenerative movement disorder, affecting 1-2% of people over 65 years of age [1]. PD is characterized by cardinal motor and non-motor symptoms, including rigidity, slowness of voluntary movements, and cognitive decline [2–4]. Intraneuronal Lewy bodies containing ubiquitinated proteins and α-synuclein are a hallmark pathologic finding in PD. The stereotypic spread of Lewy bodies in PD from the rostral brain stem to the midbrain and eventually throughout the neocortex suggests a prion-like mechanism mediating propagation of protein aggregates from neuron to neuron [5]. This temporo-spatial spread of Lewy bodies correlates with clinical progression of PD and has been replicated in several animal models [6–8]. Although much work has focused on identifying genes involved in PD, and how perturbation of these genes lead to PD pathogenesis, the mechanisms underlying PD are not yet completely understood.

Mutations in the gene *glucosidase, beta acid 1* (*GBA*), encoding the lysosomal ceramide metabolism enzyme glucocerebrosidase (GCase), are the strongest genetic risk factor for PD and dementia with Lewy bodies, increasing risk of developing PD by approximately 5-fold in *GBA* mutation carriers compared to non-carriers [9–11]. *GBA* carriers with PD are otherwise clinically similar to idiopathic PD patients, with indistinguishable response to dopaminergic medications, slightly younger age of onset by about 4 years, and higher incidence of cognitive decline [12, 13]. While some studies suggest that *GBA* carriers with PD may have a heavier burden of Lewy bodies than in non-carriers with PD, the neuropathologic features are similar [14]. Importantly, mutations in the gene *GBA* are common, having been found in 4-5% of all idiopathic PD patients [13, 15]. Recent longitudinal clinical studies have revealed that in addition to increased risk, *GBA* carriers with PD have faster progression of both motor and cognitive symptoms compared to idiopathic PD patients [16–18].

To examine how *GBA* influences PD pathogenesis, we previously developed and characterized a *Drosophila* model of *GBA* deficiency (*Gba1b*) that manifests several phenotypes reminiscent of key features of PD, including neurodegeneration, locomotor deficits, cognitive deficits, and accelerated protein aggregation in multiple tissues including the nervous system and muscle [18]. Our proteomic analysis of *Gba1b* mutant flies revealed that proteins involved in extracellular vesicle (EV) biology were dysregulated, and EVs isolated from *Gba1b Drosophila* hemolymph revealed increased levels of aggregate-prone and EV-intrinsic proteins, indicating that *GBA* deficiency alters the protein composition of EVs [19]. EVs are a heterogeneous group of membrane-bound vesicles secreted by cells that can have multiple functions, including intercellular communication through protein and nucleic acid cargo and discard of cell components outside of the cell [20, 21]. EVs can originate from the endosomal system as a result of fusion of the late endosome to the plasma membrane, releasing intraluminal vesicles as EVs into the extracellular matrix (exosomes), or from direct outward budding of the plasma membrane (microvesicles) [20, 22, 23]. EVs have been implicated in the propagation of misfolded proteins between cells in multiple neurodegenerative diseases, including PD [24–29]. α-synuclein has been found within EVs isolated from tissues of PD and dementia with Lewy bodies patients [24, 30], and *in vitro* studies have suggested that α-synuclein [31]. Accordingly, we hypothesized that tissue-specific expression of wildtype (WT) GCase might reduce the protein aggregates that accumulate in *Gba1b* mutants.

Using our *GBA*-deficient *Drosophila* model [18, 19], we examined whether GCase could be mediating propagation of protein aggregates from cell-to-cell via EVs. In this study, we found that tissue-specific expression of WT GCase in *Gba1b* mutants corrected the alterations in protein composition of *GBA*-deficient EVs, as well as protein aggregation in local and distant tissues. Perturbing two independent EV biogenesis pathways in *Gba1b* mutants resulted in tissue-specific cell-autonomous effects on protein aggregation and changes in EV protein composition but did not rescue protein aggregation in distant tissues. Finally, we observed ectopically expressed GCase in EVs, suggesting that trafficking of GCase within EVs may contribute to the observed non-cell-autonomous rescue. Our findings suggest that mutations in *GBA* result in alterations in the protein composition of EVs that promote enhanced cell-to-cell transmission of pathogenic protein aggregates. Moreover, our findings indicate that GCase can be packaged into EVs and trafficked between cells to reduce protein aggregation throughout an organism. These findings suggest a possible mechanism underlying the clinical finding that *GBA* mutation carriers not only have increased risk of developing PD, but also faster progression of disease.

## Results

### Protein aggregation in *Gba1b* mutants can be rescued cell-autonomously and non-cell-autonomously by tissue-specific *dGba1b* expression

Our prior work revealed accelerated aggregate accumulation and dysregulation of EV-related proteins in *Gba1b* mutant flies suggesting that GCase deficiency may influence cell-to-cell spread of protein aggregates. To explore whether tissue-specific *GBA* expression could reduce aggregation levels in distant tissues, we attempted to rescue the accumulated ubiquitinated protein and Ref(2)P in the brains of *Gba1b* mutants by expressing WT *Drosophila Gba1b* (*dGba1b)* in non-neural tissue. Using the *DMef-GAL4* driver to drive expression of WT *dGba1b* in muscles throughout the fly, insoluble ubiquitinated protein aggregate accumulation was reduced in both the thoraces and heads of *Gba1b* mutant flies (Fig 1A&B). This non-cell-autonomous rescue of ubiquitinated protein aggregates was dramatically apparent in whole brains (Fig 1C). However, because *DMef-GAL4* is also expressed in muscles located in the fly head that control the proboscis, we repeated this experiment using the *Act88F-GAL4* driver. *Act88F* expression is concentrated in the thoracic indirect flight muscles and is not found in the head [32]. Expression of WT *dGba1b* using the *Act88F-GAL4* driver in *Gba1b* mutants significantly reduced accumulation of insoluble ubiquitinated proteins in the thoraces and heads (Fig 1E&F) and rescued their shortened lifespan (Fig 1D).

**Fig 1.**
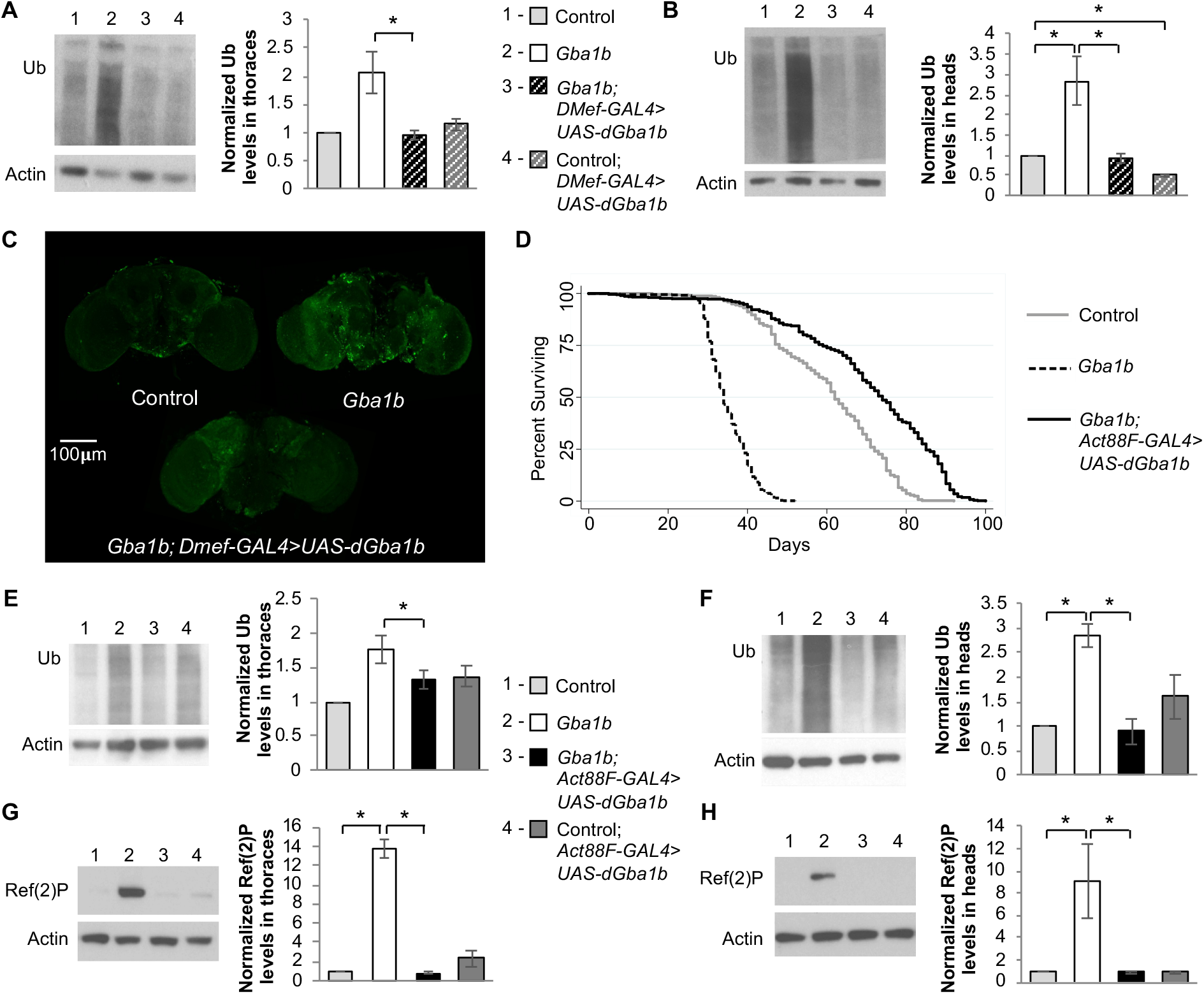
Muscle expression of *dGba1b* rescues protein aggregation and lifespan in *Gba1b* mutants. (A-C) Using the muscle-specific driver, *DMef-GAL4,* wildtype (WT) *Drosophila Gba1b* (*dGba1b)* was expressed in *Gba1b* mutants and WT revertant controls. (A,B) Homogenates were prepared from fly thoraces and heads using 1% Triton X-100 lysis buffer. Western blot analysis was performed on the Triton X-100 insoluble proteins using antibodies to ubiquitin (Ub) and Actin. Representative images and quantification of ubiquitin (Ub) in (A) thoraces and (B) heads are shown. Results are normalized to Actin loading control and control flies. (C) Representative anti-Ub immunofluorescent staining of whole brains from control, *Gba1b* mutant, and *Gba1b* mutants expressing *dGba1b* using the *DMef-GAL4* driver. (D-H) Using the indirect flight muscle specific driver, *Act88F-GAL4,* WT *dGba1b* was expressed in *Gba1b* mutants and wildtype revertant controls. (D) Kaplan-Meier survival curves of control, *Gba1b* mutants, and *Gba1b* mutants expressing WT *dGBA1b* using the *Act88F-GAL4* driver. (E-H) Homogenates were prepared from fly thoraces and heads using 1% Triton X-100 lysis buffer. Western blot analysis was performed on the Triton X-100 insoluble proteins using antibodies to ubiquitin (Ub) and Actin, and on the soluble fractions using antibodies to Ref(2)P and Actin. Representative images and quantification of ubiquitin in (E) thoraces and (F) heads and Ref(2)P in (G) thoraces and (H) heads of controls and *Gba1b* mutants with and without muscle expression of WT *dGba1b* are shown. Results are normalized to Actin in controls. At least 3 independent experiments were performed. Error bars represent SEM. *p < 0.05 by Student t-test.

Ref(2)P, the *Drosophila* ortholog of mammalian SQSTM1/p62, is important for selectively targeting ubiquitinated proteins for lysosomal degradation. While Ref(2)P is often used as a marker of autophagic flux, we have previously found it to accumulate in *Gba1b* mutants with no evidence of significantly impaired autophagy [19]. However, Ref(2)P is also required for aggregate formation under normal physiological conditions [33], and it has been shown to be released concomitantly with α-synuclein in EVs [30]; therefore we used its accumulation as a readout of protein aggregation. We anticipated that Ref(2)P accumulation in *Gba1b* mutants could also be non-cell-autonomously rescued. We indeed found that Ref(2)P accumulation was significantly decreased in both the thoraces and heads of *Gba1b* mutants expressing WT *dGba1b* in flight muscle (Fig 1G&H).

To determine whether there might be a directionality or tissue-specificity to the non-cell-autonomous rescue of protein aggregation, we expressed WT *dGba1b* in the nervous system and assessed for protein aggregates in the body. Neuronal expression of WT *dGBA1b* using the *elav*-*GAL4* driver decreased both Ref(2)P accumulation and insoluble ubiquitinated proteins in both the heads and bodies of *Gba1b* mutants (Fig 2). These results suggest that GCase may normally function to reduce the spread of protein aggregates from one cell and tissue to another.

**Fig 2.**
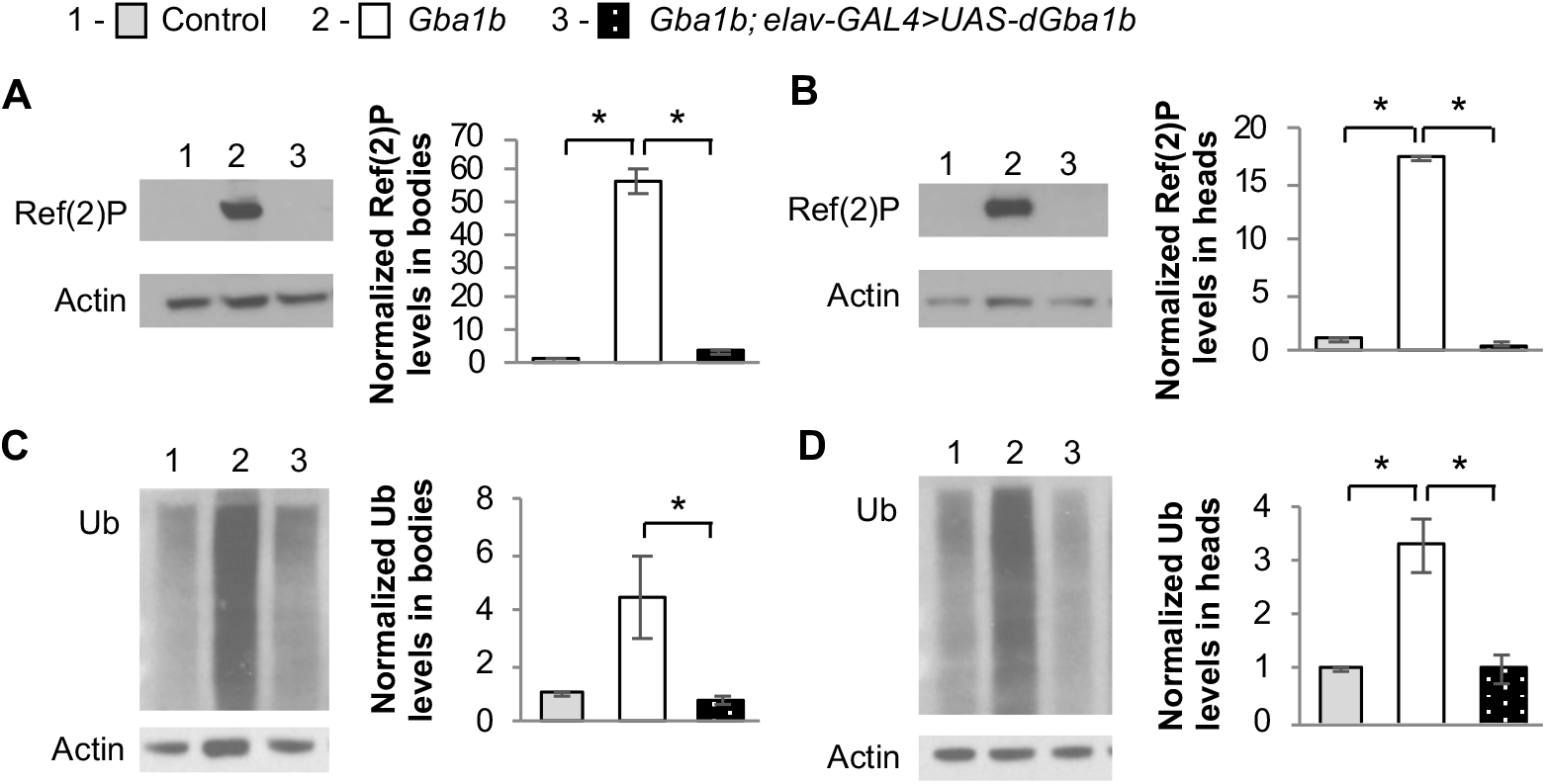
Neuronal expression of *dGba1b* rescues protein aggregation in *Gba1b* mutants. (A-D) Using the neuronal driver *elav-GAL4,* wildtype *dGba1b* was expressed in *Gba1b* mutants and wildtype revertant controls. Homogenates were prepared from fly heads and bodies using 1% Triton X-100 lysis buffer. Western blot analysis was performed on the Triton X-100 soluble fractions using antibodies to Ref(2)P and Actin and on the insoluble proteins using antibodies to ubiquitin (Ub) and Actin. Representative images and quantification of Ref(2)P in (A) bodies and (B) heads and Ub in (C) bodies and (D) heads of control and *Gba1b* mutants with and without neuronal expression of *dGba1b* are shown. Results are normalized to Actin and control. At least 3 independent experiments were performed. Error bars represent SEM. *p < 0.05 by Student t-test.

### Non-cell-autonomous rescue of protein aggregation in *Gba1b* mutants is mediated by extracellular vesicles

There are multiple mechanisms that could mediate non-cell-autonomous interactions, including exocytosis of cytoplasmic components into the extracellular matrix, direct cell-to-cell contacts, and release of cytoplasmic components via EVs. Given that our prior proteomic analysis of *Gba1b* mutants found evidence of dysregulation of EVs [19], we hypothesized that GCase influences the trafficking of aggregate-prone proteins in EVs. We isolated EVs from hemolymph and confirmed that Ref(2)P was increased in EVs from *Gba1b* mutants compared to controls (Fig 3A), as we had seen previously [19]. We also found that proteins associated with EVs, Rab11 and Rab7, were elevated (Fig 3B&C). We tested the possibility that muscle-specific expression of WT *dGba1b* would reduce the increased levels of Ref(2)P and EV-intrinsic proteins from *Gba1b* mutants. Indeed, we observed a reduction in Ref(2)P, Rab11, and Rab7 levels in EVs isolated from *Gba1b* mutants expressing WT *dGba1b* in flight muscle using the *Act88F-Gal4* driver (Fig 3A-C). However, while ubiquitinated proteins were increased in *Gba1b* mutant whole flies, they were not significantly altered in EVs isolated from *Gba1b* mutants (Fig 3D). These results indicate that expression of GCase in the muscles of *Gba1b* mutants restores normal EV content and EVs may mediate the spread of protein aggregates between tissues.

**Fig 3.**
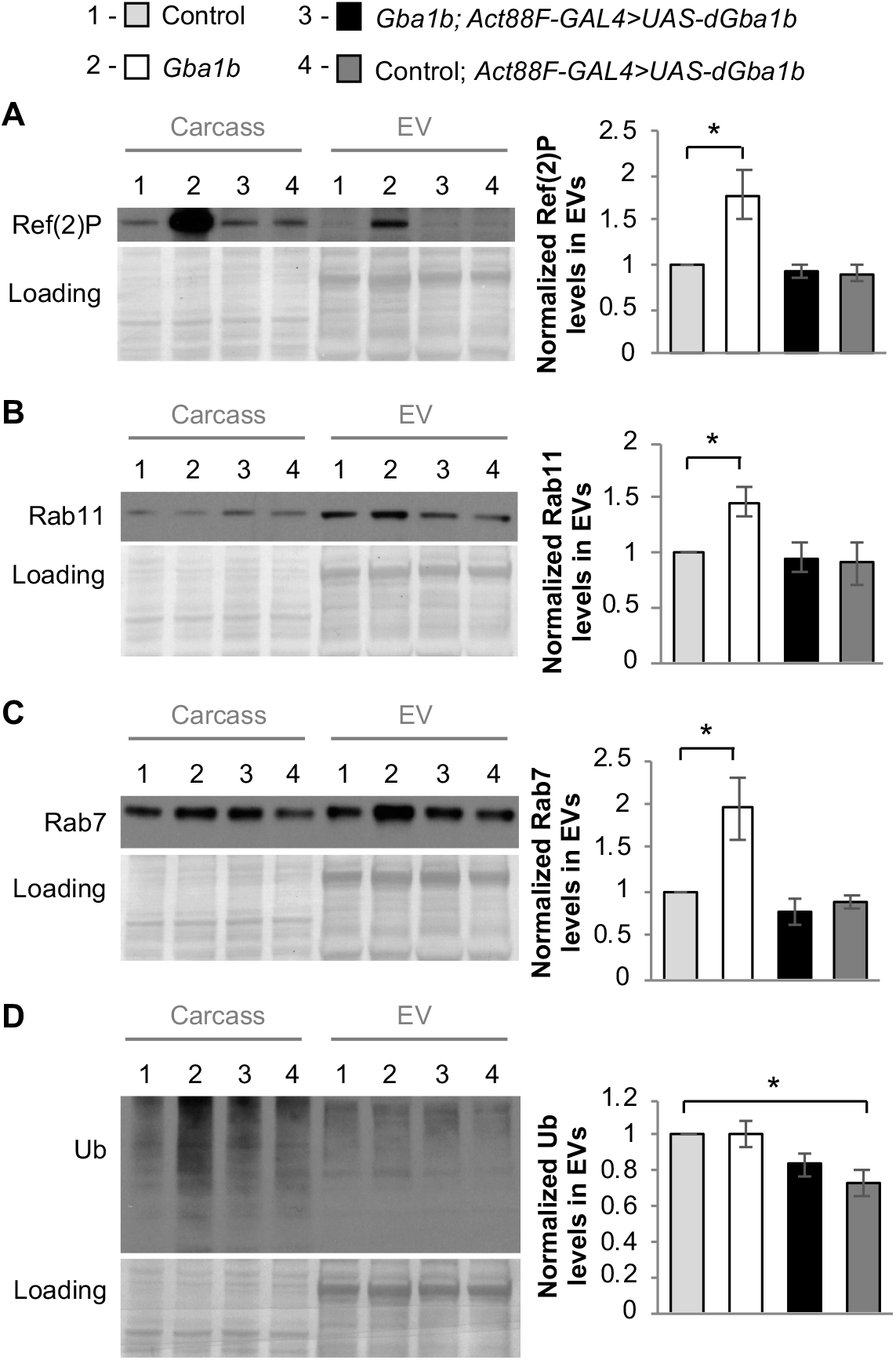
Muscle expression of *dGba1b* rescues alterations in EV-intrinsic proteins and Ref(2)P observed in EVs. (A-D) Fly carcass homogenates and isolated extracellular vesicles (EVs) from *Gba1b* mutants and wildtype revertant control flies with and without wildtype *dGba1b* expressed in indirect flight muscle using the *Act88F-GAL4* driver. Western blot analysis was performed using antibodies to (A) Ref(2)P, (B) Rab11, (C) Rab7, and (D) ubiquitin (Ub). Representative images and quantification normalized to controls are shown. Ponceau S is shown to verify equal loading. At least 3 independent experiments were performed. Error bars represent SEM. *p < 0.05 by Student t-test.

### Tissue-specific EV machinery is necessary for the spread of protein aggregation in *Gba1b* mutants

We previously found that pan-neuronal RNAi knockdown of Endosomal Sorting Complexes Required for Transport (ESCRT) proteins involved in EV biogenesis decreased the accumulation of insoluble ubiquitinated proteins and Ref(2)P cell-autonomously in the heads of *Gba1b* mutants [19]. We hypothesized that if the observed non-cell-autonomous rescue is mediated by EVs, then RNAi knockdown of ESCRT machinery might reduce protein aggregation in distant tissues in *Gba1b* mutants by decreasing production of dysregulated EVs that promote propagation of protein aggregates. We confirmed that neuronal RNAi knockdown of *Multivesicular body subunit 12 (Mvb12)*, a component of the ESCRT machinery required for EV biogenesis [34, 35], cell-autonomously decreased ubiquitinated proteins and Ref(2)P levels in heads (Fig 4A&B). However, RNAi knockdown of *Mvb12* in muscle did not rescue increased ubiquitinated proteins or Ref(2)P levels in the heads of *Gba1b* mutants (Fig 4C&D). This suggests that knockdown of ESCRT-dependent EV biogenesis may be specifically important for the cell-autonomous regulation of protein aggregation in neurons but is not involved in the non-cell-autonomous rescue of brain protein aggregation by muscle-specific *dGba1b* expression.

**Fig 4.**
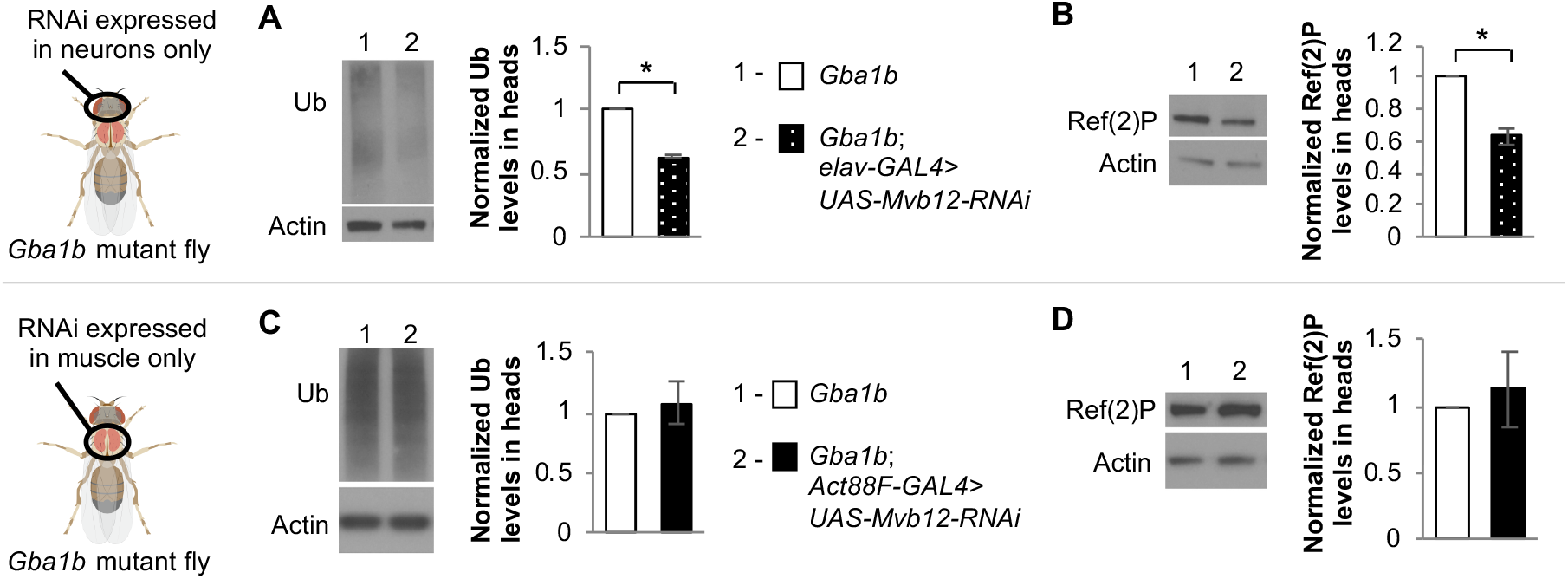
Tissue-specific knockdown of *Mvb12* in *Gba1b* mutants differentially alters ubiquitinated proteins and Ref(2)P. (A-D) *Multivesicular body sorting factor 12* (*Mvb12)-*RNAi was expressed using tissue-specific drivers in *Gba1b* mutants. Homogenates were prepared from fly heads using 1% Triton X-100 lysis buffer. Western blot analysis was performed on the Triton X-100 insoluble proteins using antibodies to ubiquitin (Ub) and Actin, and on the soluble fractions using antibodies to Ref(2)P and Actin. (A-B) Representative images and quantification of (A) Ub and (B) Ref(2)P in the heads of flies with and without *Mvb12* knockdown in neurons using the *elav-GAL4* driver. (C-D) Representative images and quantification of (C) Ub and (D) Ref(2)P in the heads of flies with and without *Mvb12* knockdown in flight muscle using the *Act88F-GAL4* driver. Results are normalized to *Gba1b* mutants without RNAi expression. At least 3 independent experiments were performed. Error bars represent SEM. *p < 0.05 by Student t-test.

We next examined knockdown of *neutral sphingomyelinase* (*nSMase*), a lipid-modifying enzyme important for the formation and release of EVs [36, 37]. nSMase hydrolyzes sphingomyelin, producing phosphocholine and ceramide. This enzyme has been implicated in multiple cellular functions, including inflammatory responses, reaction to lung and cardiac pathology, synaptic regulation, and release of EVs independent of ESCRT machinery [38]. We again hypothesized that tissue-specific RNAi knockdown of *nSMase* in *Gba1b* mutants could reduce protein aggregation in distant tissues by reducing the production of dysregulated EVs promoting protein aggregation. Instead, we observed increased accumulation of ubiquitinated proteins and Ref(2)P in the thoraces and a trend towards an increase in ubiquitinated proteins and Ref(2)P in the heads of *Gba1b* mutants after knockdown of *nSMase* expression in flight muscle (Fig 5A-D). However, knocking down *nSMase* in neuronal tissue did not change Ref(2)P or ubiquitinated protein levels in the heads of *Gba1b* mutants (Fig 5E&F). These results suggest that nSMase-mediated release of EVs may be important for the cell-autonomous regulation of protein aggregation in muscles but not neurons. We next examined whether EV cargo is altered in *Gba1b* mutants with RNAi knockdown of *nSMase* in muscle. We found increased Ref(2)P and ubiquitinated protein levels in EVs isolated from *Gba1b* mutants with *nSMase* knocked down in muscle, but no change in the levels of Rab11 or Rab7, which are already increased in EVs from *Gba1b* mutants (Fig 6). This suggests that nSMase may regulate the protein cargo of EVs but may not alter the abundance of EVs.

**Fig 5.**
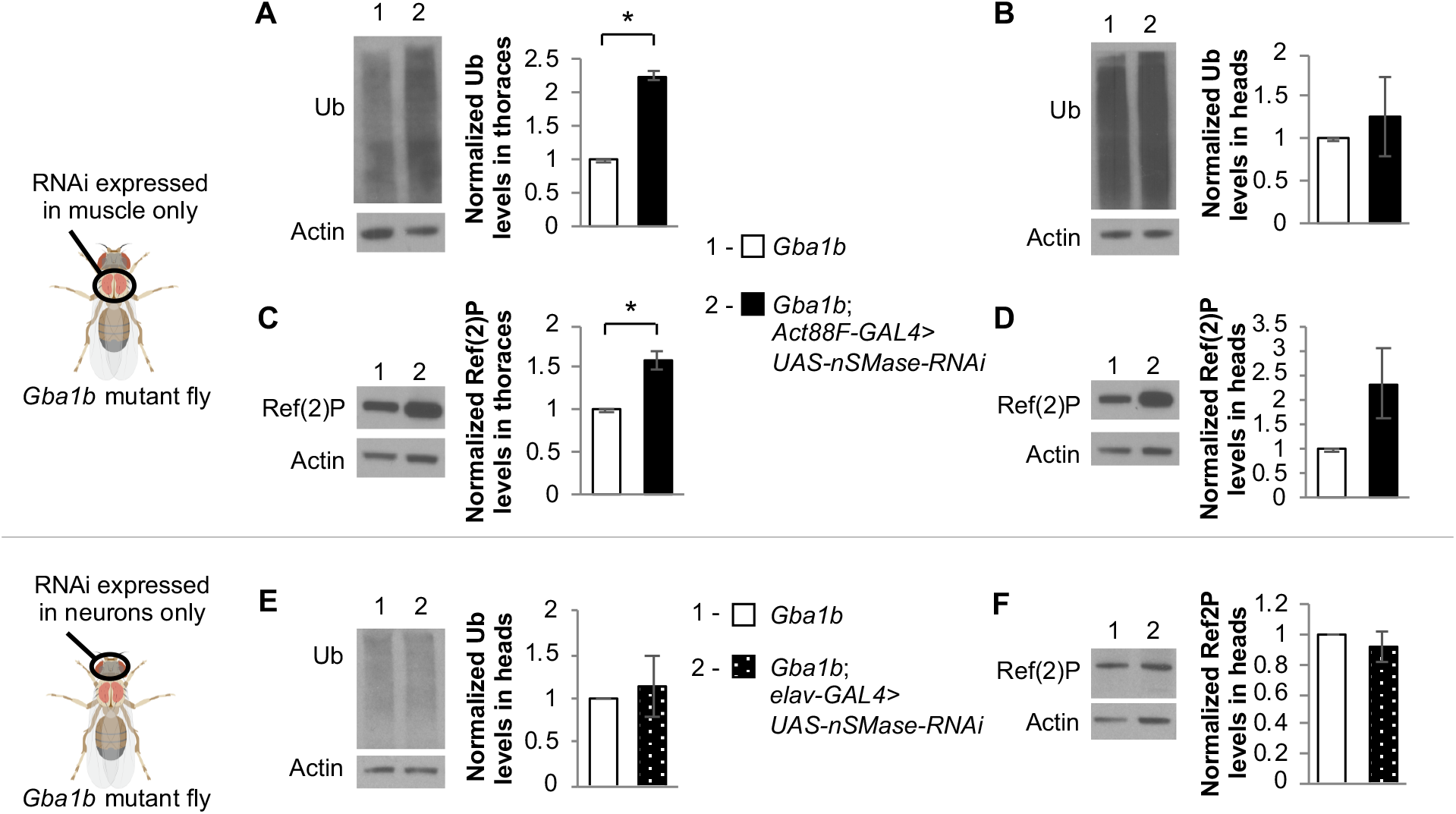
Tissue-specific knockdown of nSMase in *Gba1b* mutants differentially alters ubiquitinated proteins and Ref(2)P. (A-F) *Neutral sphingomyelinase (nSMase)-*RNAi was expressed using tissue-specific drivers in *Gba1b* mutants. Homogenates were prepared from fly heads and thoraces using 1% Triton X-100 lysis buffer. Western blot analysis was performed on the Triton X-100 insoluble proteins using antibodies to ubiquitin (Ub) and Actin, and on the soluble fractions using antibodies to Ref(2)P and Actin. (A-D) Representative images and quantification of ubiquitin in (A) thoraces and (B) heads and Ref(2)P in (C) thoraces and (D) heads of flies with and without nSMase knockdown in flight muscle using the *Act88F-GAL4* driver. (E,F) Representative images and quantification of (E) ubiquitin and (F) Ref(2)P in the heads of flies with and without nSMase knockdown in neurons using the *elav-GAL4* driver. Results are normalized to *Gba1b* mutants without RNAi expression. At least 3 independent experiments were performed. Error bars represent SEM. *p < 0.05 by Student t-test.

**Fig 6.**
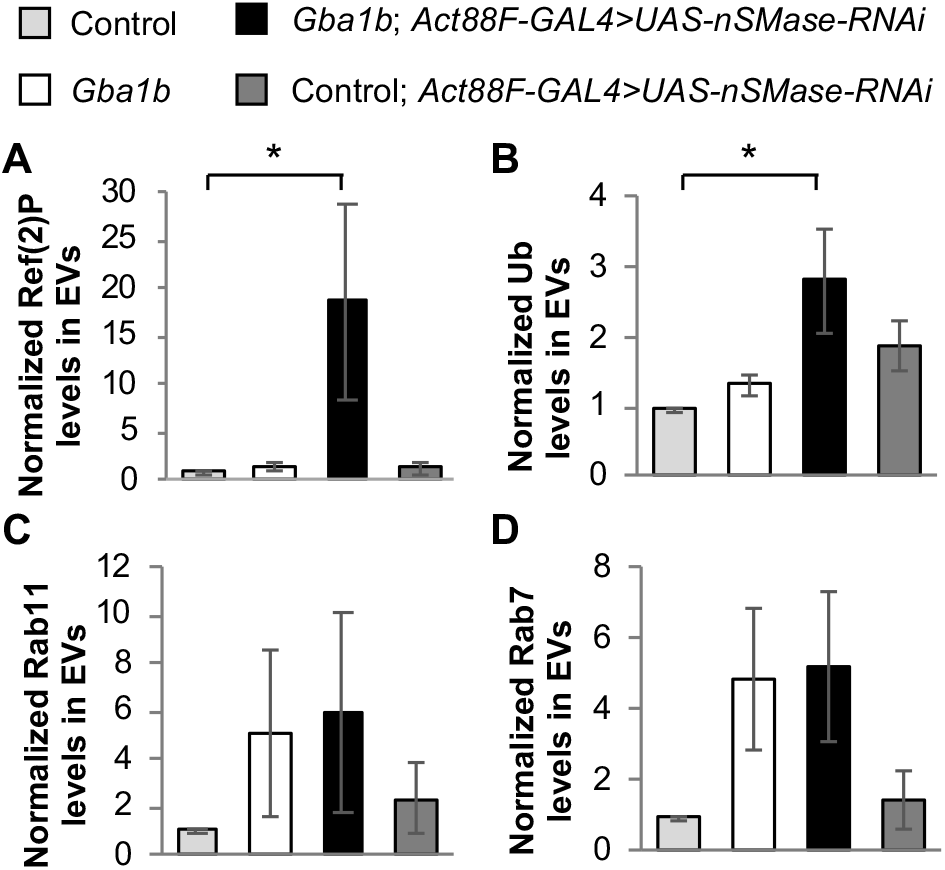
Muscle-specific knockdown of nSMase alters EV protein cargo. (A-D) *Neutral sphingomyelinase (nSMase)-*RNAi was expressed using the flight muscle driver *Act88F-GAL4* in *Gba1b* mutants and wildtype revertant controls. Isolated EVs from these flies were prepared in RIPA buffer. Quantification of western blot analysis of (A) Ref(2)P, (B) ubiquitin, (C) Rab11, and (D) Rab7 in the EV fraction are shown. Results are normalized to control. At least 3 independent experiments were performed. Error bars represent SEM. *p < 0.05 by Student t-test.

### Ectopically expressed GCase is trafficked within extracellular vesicles

Our finding that *GBA* expression in muscle rescues the protein composition in EVs indicates that non-cell-autonomous rescue is mediated at least in part by preventing spread of aggregated proteins trafficked by EVs. However, EV-mediated trafficking of GCase could also be another contributing factor to the non-cell-autonomous rescue of protein aggregation in *Gba1b* mutants. To test this possibility, we investigated whether ectopically expressed GCase travels to distant tissues via EVs. Because there are no antisera that recognize *Drosophila* GCase, we examined ectopic expression of human *GBA (hGBA)* in *Gba1b* mutants. We previously found ubiquitous expression of WT *hGBA* to be sufficient to partially rescue the shortened lifespan of *Gba1b* mutants [18]. Here, we found that expression of WT *hGBA* in flight muscle also reduced ubiquitinated protein aggregation and Ref(2)P accumulation in *Gba1b* mutants in both thoraces and heads (Fig 7A-D). EVs isolated from *Gba1b* mutants expressing *hGBA* in flight muscle were found to have decreased Rab11 levels compared to controls (Fig 8A&B), confirming that human GCase (hGCase) can also revert alterations in EVs due to *dGba1b* deficiency. These results support a functional equivalence between *Drosophila* and hGCase in reducing the formation of protein aggregates. Interestingly, we were also able to detect hGCase in both the thoraces and heads of *Gba1b* mutants expressing *hGBA* in flight muscle (Fig 7E), indicating that hGCase itself is able to travel to distant tissues. Furthermore, we detected hGCase in EVs isolated from hemolymph of flies expressing *hGBA* in muscle (Fig 8A). To determine whether hGCase is localized within EVs or associated on the outer surface of EVs, we added Proteinase K to the isolated EV fraction. We found hGCase to be resistant to Proteinase K digestion of the EV fraction but degraded if EVs were first treated with a detergent to disrupt vesicular membranes before Proteinase K digestion (Fig 8C). These results indicate that GCase can be incorporated into EV cargo, and that WT GCase may be trafficked by EVs to distant tissues to reduce protein aggregation.

**Fig 7.**
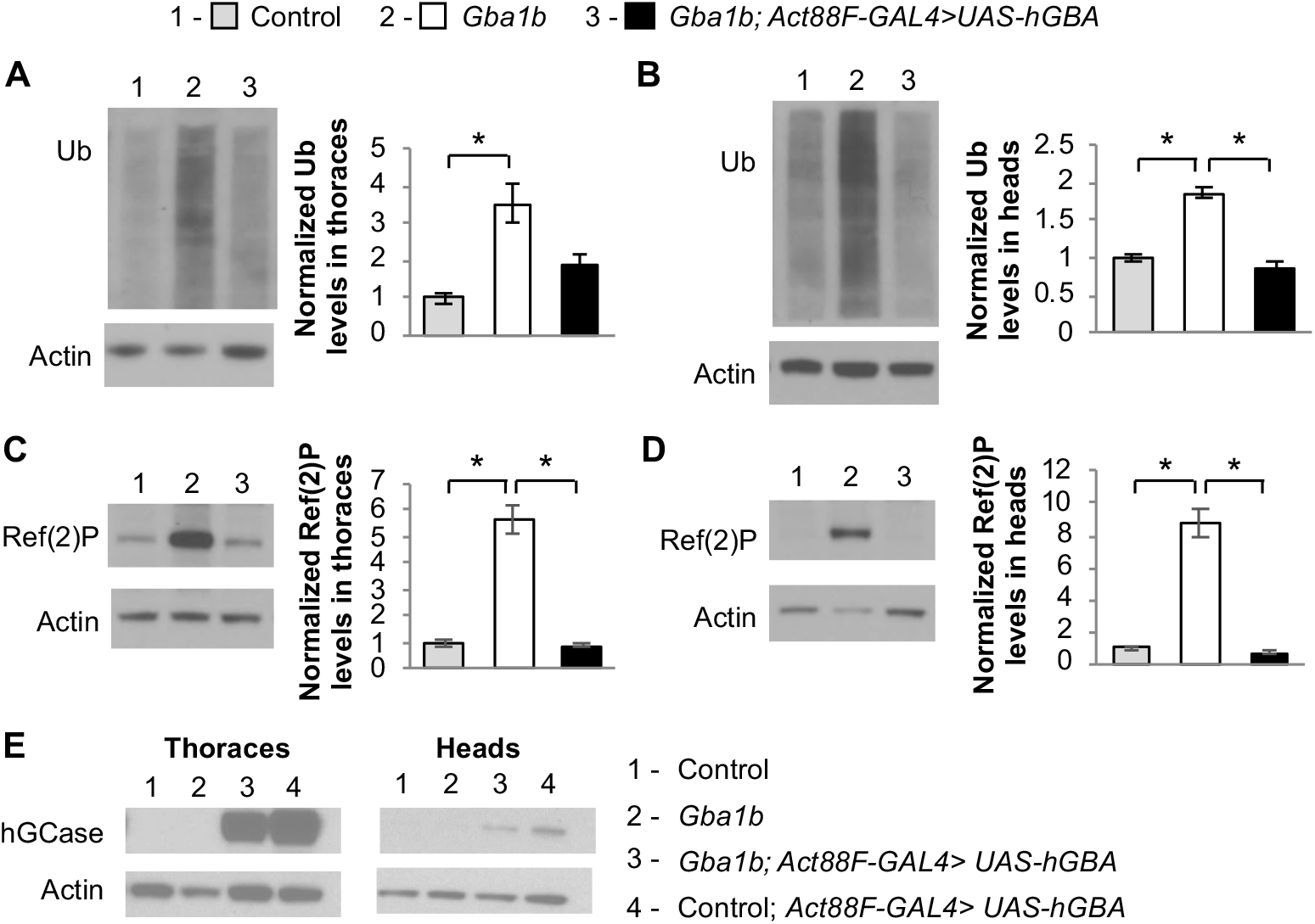
Muscle expression of human *GBA* suppresses protein aggregation in *Gba1b* mutants. (A-E) Using the flight muscle specific driver, *Act88F-GAL4,* wildtype (WT) human *GBA* (*hGBA*) was expressed in *Gba1b* mutant and WT revertant controls. Homogenates were prepared from fly heads and thoraces using 1% Triton X-100 lysis buffer. Western blot analysis was performed on the Triton X-100 insoluble proteins using antibodies to ubiquitin (Ub) and Actin, and on the soluble fractions using antibodies to Ref(2)P, Actin, and human glucocerebrosidase (hGCase). Representative images and quantification of ubiquitin in (A) thoraces and (B) heads and Ref(2)P in (C) thoraces and (D) heads of *Gba1b* mutant flies with and without muscle expression of WT *hGBA* are shown. Results are normalized to Actin and control. At least 3 independent experiments were performed. Error bars represent SEM. *p < 0.05 by Student t-test. (E) Antibody detecting hGCase in the thoraces and heads of control and *Gba1b* mutant flies with and without muscle expression of WT *hGBA*.

**Fig 8.**
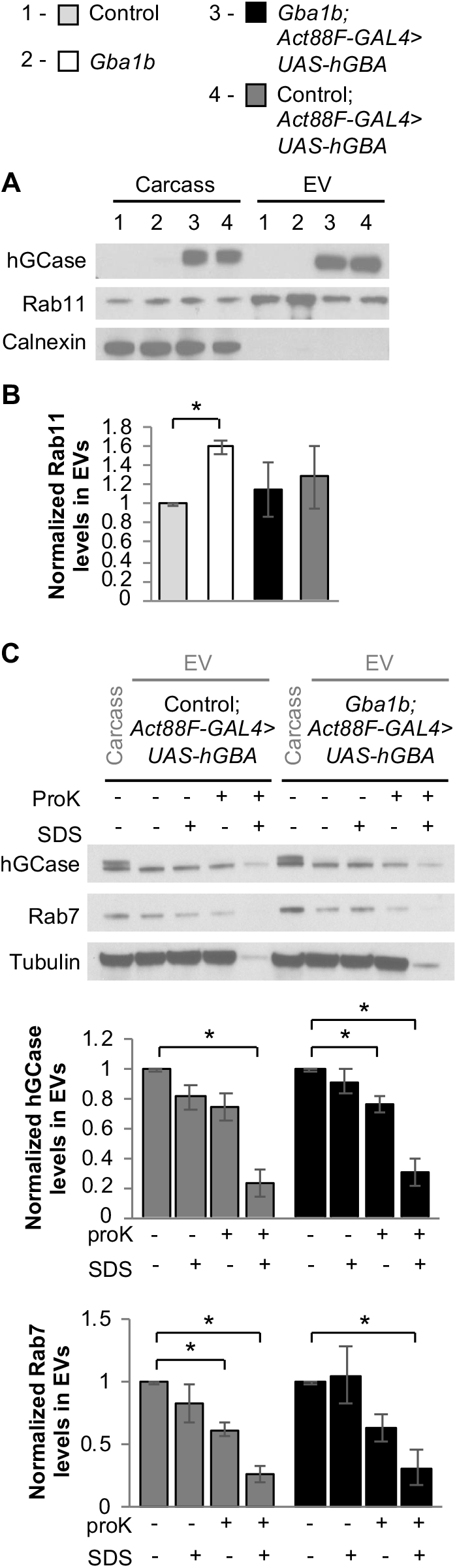
Ectopically expressed human GCase is trafficked within EVs. (A-B) Fly carcass homogenates and isolated extracellular vesicles (EVs) from *Gba1b* mutants and wildtype (WT) revertant controls with and without WT human *GBA* (*hGBA)* expressed in flight muscle using the *Act88F-GAL4* driver were prepared using RIPA buffer. Western blot analysis was performed using antibodies to human glucocerebrosidase (hGCase), Rab11, Rab7, and Tubulin. (A) Representative images of hGCase and Rab11 in carcass and isolated EVs from control and *Gba1b* mutant flies with and without muscle expression of *hGBA* are shown. The blots were also probed with antibodies to Calnexin (Cnx99A) to confirm the purity of the EV samples. (B) Levels of Rab11 in the EV fraction are quantified and normalized to controls. (C) Representative images of hGCase, Rab7, and Tubulin in carcass and isolated EVs from control and *Gba1b* mutant flies with muscle expression of WT *hGBA* with and without Proteinase K (ProK) and/or SDS exposure. Levels of hGCase and Rab7 in the EV fraction are quantified. Results are normalized to EVs without ProK or SDS exposure. At least 3 independent experiments were performed. Error bars represent SEM. *p < 0.05 by Student t-test.

## Discussion

Many genetic influences of PD have now been identified, and much work has been focused on how these genes lead to protein aggregation through mechanisms such as protein misfolding and autophagy defects. However, none of these genes have been implicated in cell-to-cell spread of pathogenic protein aggregates, which closely correlates with clinical disease progression. Our proteomic analysis and non-cell-autonomous rescue of protein aggregation in *Gba1b* mutants has led us to hypothesize that *GBA* mutations may influence the rate of propagation of protein aggregates between neurons. Our work suggests a link between *GBA* mutations and faster spread of intracellular protein aggregates via a novel EV-mediated mechanism, possibly explaining the recent clinical finding that *GBA* mutations accelerate the progression of clinical disease. Using a *Drosophila* model of *GBA* deficiency that manifests accelerated protein aggregation, we found that expressing WT GCase in specific tissues of a *GBA*-deficient fly can not only rescue protein aggregation cell-autonomously and in distant tissues, but also rescue alterations in protein cargo observed in EVs isolated from *Gba1b* mutant hemolymph. Interestingly, ectopically expressed WT GCase itself was found within EVs of *GBA*-deficient flies, suggesting that the non-cell-autonomous rescue due to GCase expression is mediated by both reduction in aggregated proteins in EVs and trafficking of GCase via EVs to distant cells and tissues. Perturbing EV biogenesis through decreased expression of ESCRT machinery or ESCRT-independent nSMase affected protein aggregation in local tissues in a tissue-dependent manner. Together, these findings suggest that mutations in *GBA* result in the accelerated spread of protein aggregates through dysregulation of proteins trafficked by EVs.

Although our model of *GBA* mutations promoting spread of protein aggregates via EVs is novel, the idea that proteostasis can be maintained in a non-cell-autonomous fashion is well supported in the literature. For example, in *C. elegans*, misfolded α-synuclein accumulating in endo-lysosomal vesicles was found to be transmitted from muscle to the hypodermis, a nearby tissue, for degradation [39]. It is possible that a non-cell-autonomous mechanism is necessary because certain tissues may be more efficient in reducing protein aggregation. This has been previously described, where overexpression of *FOXO* in *Drosophila* muscle decreased aging-related protein aggregates in muscle as well as brain and other distant tissues, but *FOXO* overexpression in adipose tissue was unable to prevent protein aggregation in muscle [40]. In our model, overexpressing *dGba1b* in *Drosophila* muscle or neuronal tissue prevented accumulation of protein aggregates throughout the organism, however overexpression of WT GCase in midgut and fat body did not significantly reduce protein aggregation in the brain (S Fig 1). These discrepancies could be due to tissue-specific differences influencing biogenesis of EVs, such as metabolic rate or endovesicular trafficking intrinsic to specific cells.

Our unexpected results from perturbations of EV biogenesis machinery suggest that the EV-mediated regulation of protein aggregation is tissue-specific and complex. Because an increase in EV-intrinsic proteins and alteration of protein cargo were observed in *Gba1b* mutants [19], we anticipated that genetic perturbations decreasing the biogenesis of EVs might also rescue protein aggregation non-cell-autonomously by reducing the production of dysregulated EVs. However, decreased expression of ESCRT component *Mvb12* or ESCRT-independent *nSMase* in muscle did not rescue protein aggregation in heads, suggesting that a tissue-specific decrease in biogenesis of dysregulated EVs is not sufficient to reduce protein aggregation in the rest of the organism, and the cargo of EVs may need to be corrected to reduce spread of protein aggregation. A possible explanation for why decreased muscle expression of *nSMase* enhanced cell-autonomous protein aggregation and EV protein cargo alterations is that both GCase and nSMase enzymatically produce ceramide. If GCase-deficient phenotypes are dependent on a relative reduction in ceramide, decreased *nSMase* expression could exacerbate *Gba1b* mutant phenotypes. In contrast, decreased expression of *nSMase* in the nervous system had no effect on protein aggregation in the head and neuronal knockdown of *Mvb12* cell-autonomously rescued protein aggregation. These discrepancies between perturbation of EV biogenesis pathways and tissue types could be due to cell-specific compensatory mechanisms or intrinsic metabolic demands and solicit further investigation.

Our work suggests that GCase deficiency influences EV biogenesis to promote faster propagation of pathogenic protein aggregates throughout the tissues of an organism, which may be a compensatory response to cell-autonomous lysosomal stress. In the initial characterization of our *Drosophila GBA*-deficient model we found accelerated insoluble ubiquitinated protein aggregates, accumulation of Ref(2)P, and oligomerization of ectopically expressed human α-synuclein in *Gba1b* mutants, suggesting an impairment in lysosomal degradation [18, 42]. A similar *GBA*-deficient *Drosophila* model also found evidence of lysosomal dysfunction, including enlarged lysosomes in *GBA*-deficient brains [43]. However, our proteomic analysis of *Gba1b* mutants did not support a profound impairment in autophagy, but instead suggested dysregulation of EVs with altered protein cargo which could be suppressed locally with knockdown of genes encoding ESCRT machinery important for EV biogenesis [19]. Based on these results, we believe that our initial observations of increased insoluble ubiquitinated proteins and Ref(2)P in *Gba1b* mutants are due to lysosomal stress. One possible explanation for our proteomic findings is that there may be a compensatory increase in EV biogenesis and packaging of autophagy substrates within EVs for discard outside of the cell in *Gba1b* mutants. Such an increase may have prevented us from detecting defects in autophagy. Upregulation of EV biogenesis may be cell-autonomously neuroprotective in the setting of lysosomal stress, particularly in aggregation-prone neurodegenerative diseases such as PD [42]. It was recently demonstrated in a human neuronal cell culture model of PD that inhibiting macroautophagy protects against α-synuclein-induced cell death by promoting the release of α-synuclein-containing EVs [44]. However, it remains possible that upregulating EV biogenesis may relieve lysosomal stress within cells containing aggregate-prone proteins, while simultaneously promoting the spread of protein aggregates between cells and throughout the organism.

Our work suggests a novel mechanism for *GBA* in reducing the spread of pathogenic protein aggregation from cell-to-cell via regulation of EV protein cargo, but many key questions remain. To better understand the progression of neurodegenerative diseases, we must uncover the mechanisms by which GCase deficiency alters EV protein content and biogenesis, identify the specific changes in EVs facilitating propagation of pathogenic protein aggregates, and determine how these changes influence recipient cells internalizing dysregulated EVs. GCase is a critical enzyme in ceramide metabolism, hydrolyzing glucosylceramide into glucose and ceramide. Ceramides are a key component of EV membranes, and alterations in ceramide metabolism due to GCase deficiency may directly influence EV biogenesis and protein cargo trafficked via EVs. Further studies using this *Drosophila* model and mammalian cell culture models should better elucidate how GCase deficiency alters the protein cargo of EVs to reduce propagation of pathogenic protein aggregates, as well as whether endogenous GCase is enzymatically functional when trafficked to distant tissues via EVs. Understanding this mechanism could have broad implications in understanding the pathogenesis of aggregate-prone neurodegenerative diseases and reveal new therapeutic targets to slow or halt disease progression.

## Materials and methods

### *Drosophila* strains and culture

Fly stocks were maintained on standard cornmeal-molasses food at 25°C. The *Gba1b* homozygous null mutant (*Gba1b*^*ΔTT*^), isogenic control (*Gba1b^+^)*, and *UAS-dGba1b* alleles have been previously described.[18] All other strains and alleles were obtained from the Bloomington Stock Center: *elav-GAL4* (458); *Act88F-GAL4* (38459); *DMef-GAL4* (27390); *UAS-Mvb12-RNAi* (43152); *UAS-nSMase-RNAi* (36759); *Gba1b^MB03039^(23602)*. In Figures 6 & 7, we used the following genotypes for the experiments involving the *nSMase* RNAi transgene: control = *Gba1b^+^/Gba1b^MB03039^*; *Gba1b* mutant = *Gba1b*^*ΔTT*^/*Gba1b^MB03039^*. This combination of *Gba1b* mutant alleles, which we used for ease of recombination with the transgene, produce the same biochemical abnormalities found in *Gba1b*^*ΔTT*^ homozygotes [19].

### Lifespan analysis

Longevity assays were conducted at 25°C. Groups of 10–20 age-matched flies were collected at 0–24 hours old and transferred to fresh standard food every 2–3 days. The number of dead flies was recorded during each transfer. Transfers were continued until all flies died. Kaplan-Meier lifespan curves were generated using Stata (StataCorp, College Station, TX), and analyzed by Cox proportional hazard models for statistically significant differences in survival between tested genotypes.

### Preparation of Triton-soluble and insoluble fractions

Tissues from 10-day-old flies (6 females and 6 males per sample) were homogenized in Triton lysis buffer (50 mM Tris-HCl pH 7.4, 1% Triton X-100, 150 mM NaCl, 1 mM EDTA), and then spun at 15,000 x *g* for 20 min. The detergent-soluble supernatant was collected and mixed with an equal volume of 2x Laemmli buffer (4% SDS, 20% glycerol, 120 mM Tris-Cl pH 6.8, 0.02% bromophenol blue, 2% β-mercaptoethanol), and the same buffer was used to resuspend the Triton-insoluble pellet. All samples were boiled for 10 minutes. The Triton-insoluble protein extracts were then cleared of debris by centrifugation at 15,000 x *g* for 10 minutes, followed by collection of the supernatant. At least three independent experiments were performed.

### Western blotting

Proteins were separated by SDS-PAGE on 4%-20% MOPS-acrylamide gels (GenScript Express Plus, M42012) and electrophoretically transferred onto Immobilon PVDF membranes (Fisher, IPVH00010). Immunodetection was performed using the iBind Flex Western Device (Thermo Fisher, SLF2000). Antibodies were used at the following concentrations: 1:10,000 mouse anti-Actin (Chemicon/Bioscience Research Reagents, MAB1501), 1:250 mouse anti-Rab11 (BD Transduction Laboratories, 610657), 1:50 mouse anti-Rab7 (DSHB Rab7), 1:200 rabbit anti-Ref(2)P (Abcam, ab178440), 1:500 mouse anti-ubiquitin (Santa Cruz, sc-8017), 1:10,000 mouse anti-tubulin (DSHB 12G10), 1:1500 rabbit anti-human GBA1 (Sigma, G4171), 1:800 mouse anti-Cnx99A (DSHB, Cnx99A 6-2-1). HRP secondary antibodies were used as follows: 1:500 to 1:1000 anti-mouse (BioRad, 170–6516) and 1:500 to 1:1000 anti-rabbit (BioRad, 172–1019). Signal was detected using Pierce ECL Western Blotting Substrate (Fisher, 32106). Densitometry measurements of the western blot images were performed using Fiji software [45]. For homogenates, signal was normalized either to Actin or Ponceau S [46, 47]. Normalized western blot data were log-transformed when necessary to stabilize variance before means were compared using Student *t*-test. Each experiment was performed at least three times.

### Immunohistochemistry

10-day-old adult brains were dissected in cold Schneider’s *Drosophila* medium (Thermo Fisher, 21720), and fixed in 4% paraformaldehyde/PBS for 30 min. Samples were washed in 0.1% Triton X-100/PBS. Fixed brains were stained with 1:200 mouse anti-poly Ubiquitin FK2 (Enzo, BML-PW8810), then anti-mouse Alexa 488 (1:200) and were mounted using ProLong Gold anti-fade medium (Molecular Probes, P10144).

### Extraction of hemolymph and preparation of EV fractions

Hemolymph was obtained from 30 flies (15 males and 15 females, 10 to 11 days old) per sample. All flies were frozen with liquid nitrogen and decapitated by vortexing. Frozen flies were placed in a 1.7-mL tube containing a volume of PBS scaled to the number of flies used (2 μL/fly) and thawed for 5 min at room temperature. The tubes were then centrifuged at 5000 x *g* for 5 min at 4°C, after which the extracted hemolymph (supernatant) was centrifuged for 30 min at 10,000 x *g* at 4°C to remove cell debris and the cell-free supernatant was collected. The supernatant was then filtered with Ultrafree 0.22 μm spin filters (Fisher, UFC30GV0S) and centrifuged at 3000 x *g* for 5 min at 4°C; this was the EV fraction. In order to obtain whole-fly homogenate from the same animals used for collection of hemolymph, the bodies from four flies (2 males and 2 females) were homogenized in RIPA buffer (150mM NaCl, 1% Nonidet P-40, 0.5% Sodium deoxycholate, 0.1% SDS, 50mM Tris pH 8, diluted 1:1 with ddH_2_0), centrifuged at 10,000 x *g* at 4°C for 5 min, and then the supernatant was transferred to a new tube. An equal volume of 2x Laemmli buffer (4% SDS, 20% glycerol, 120 mM Tris-Cl pH 6.8, 0.02% bromophenol blue, 2% β-mercaptoethanol) was added to the EV fractions and also to the whole-fly protein homogenates, and all samples were boiled for 10 min and then stored at −80°C. The experiment was repeated at least three times.

To determine if proteins were contained within EVs, before adding the Laemmli buffer, the EV fractions were incubated with PBS for 5 min at room temperature and then 0.5 μg/μL Proteinase K was added for 30 min at 4°C to digest external proteins or with 1% sodium dodecyl sulfate (SDS) for 5 min at room temperature and then 0.5 μg/μL Proteinase K was added for 30 min at 4°C to disrupt vesicular membranes and digest proteins. After incubation, Laemmli buffer was added and samples were boiled preparing them for western blotting.

## Acknowledgements

Kim Miller (Digital Microscopy Center, Virginia Merrill Bloedel Hearing Research Center, University of Washington) for technical assistance with confocal imaging and analysis; Biorender for *Drosophila* icon figures; Evelyn S. Vincow, Colby L. Samstag, Bernice Lin, and all members of the Pallanck and Davis labs for critical review and discussion of this work and manuscript.

## Supporting information

**S1 Fig.**
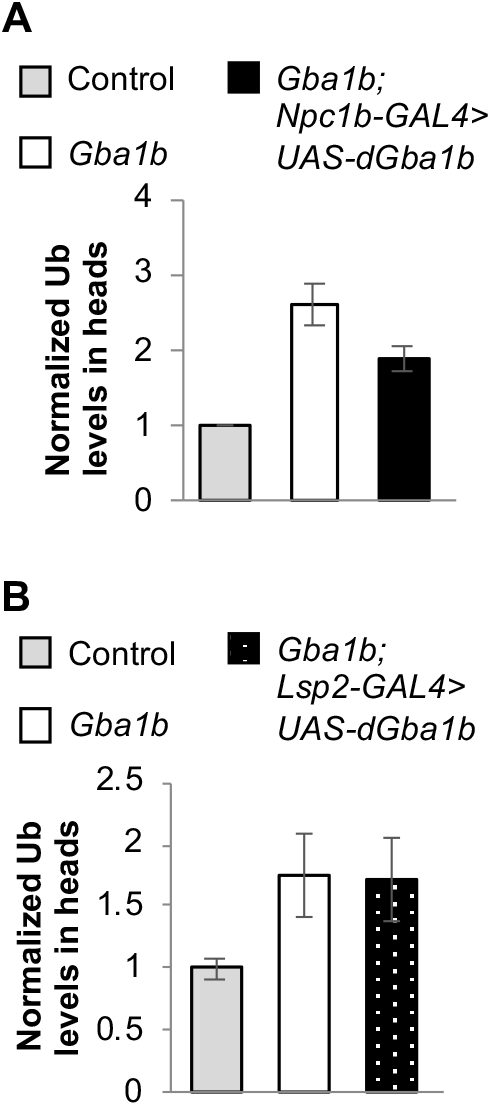
Midgut and fat body expression of *dGba1b* does not rescue protein aggregation in *Gba1b* mutants. (A) Using the midgut driver *Npc1b-GAL4,* wildtype *dGba1b* was expressed in *Gba1b* mutants and wildtype revertant controls. Homogenates were prepared from fly heads using 1% Triton X-100 lysis buffer. Western blot analysis was performed on the Triton X-100 insoluble proteins using antibodies to ubiquitin (Ub) and Actin. Quantification of Ub in (A) heads of control and *Gba1b* mutants with and without midgut expression of *dGba1b* are shown. (B) Quantification of Ub in the heads of flies with and without *dGba1b* expression in fat body using the *Lsp2-GAL4* driver. Results are normalized to Actin and control. At least 3 independent experiments were performed. Error bars represent SEM. *p < 0.05 by Student t-test.

